# Cortical patterns shift from sequence feature separation during planning to integration during motor execution

**DOI:** 10.1101/2022.07.13.499902

**Authors:** Rhys Yewbrey, Myrto Mantziara, Katja Kornysheva

**Author notes:** **Correspondence:** Dr Katja Kornysheva at, Centre for Human Brain Health, School of Psychology, University of Birmingham, Birmingham, B15 2TT, UK. **Author contributions:** R.Y., M.M. and K.K. conceived the experiments; R.Y., and K.K. formulated the hypotheses; R.Y., M.M. and K.K. collected the data; R.Y. and K.K. designed the analysis; R.Y. and K.K. performed the analyses; R.Y. and K.K. wrote the original version of the manuscript; All authors contributed to editing of the manuscript.

## Abstract

Performing sequences of movements from memory and adapting them to changing task demands is a hallmark of skilled human behaviour, from handwriting to playing a musical instrument. Prior studies showed a fine-grained tuning of cortical primary motor, premotor, and parietal regions to motor sequences – from the low-level specification of individual movements to high-level sequence features like sequence order and timing. However, it is not known how tuning in these regions unfolds dynamically across planning and execution. To address this, we trained 24 healthy right-handed participants to produce four five-element finger press sequences with a particular finger order and timing structure in a delayed sequence production paradigm entirely from memory. Local cortical fMRI patterns during preparation and production phases were extracted from separate ‘No-Go’ and ‘Go’ trials, respectively, to tease out activity related to these peri-movement phases. During sequence planning, premotor and parietal areas increased tuning to movement order and timing, irrespective of their combinations. In contrast, patterns reflecting the unique integration of sequence features emerged in these regions during execution only, alongside timing-specific tuning in the ventral premotor, supplementary motor, and superior parietal areas. This was in line with the participants’ behavioural transfer of trained timing, but not of order to new sequence feature combinations. Our findings suggest a general neural state shift from high-level feature separation to low-level feature integration within cortical regions for movement execution. Recompiling sequence features trial-by-trial during planning may enable flexible last-minute adjustment before movement initiation.

## Introduction

Skilled sequences of movements performed from memory are regarded as a hallmark of human dexterity ^1–3^. They are essential building blocks of everyday skilled behaviours, from typing and handwriting, to tying shoelaces, or playing a musical instrument (Figure 1a). In addition to the order of movements in a sequence, the temporal accuracy of the movements can be crucial to the success of the task, such as when typing a Morse code, or performing a martial arts sequence. While our knowledge of the neural control of motor sequences in the primate brain, particularly across cortical regions, has significantly expanded over the last decade, it is not well understood how higher-order features of sequences such as movement order and timing are integrated trial-by-trial for skilled performance.

**Figure 1.**
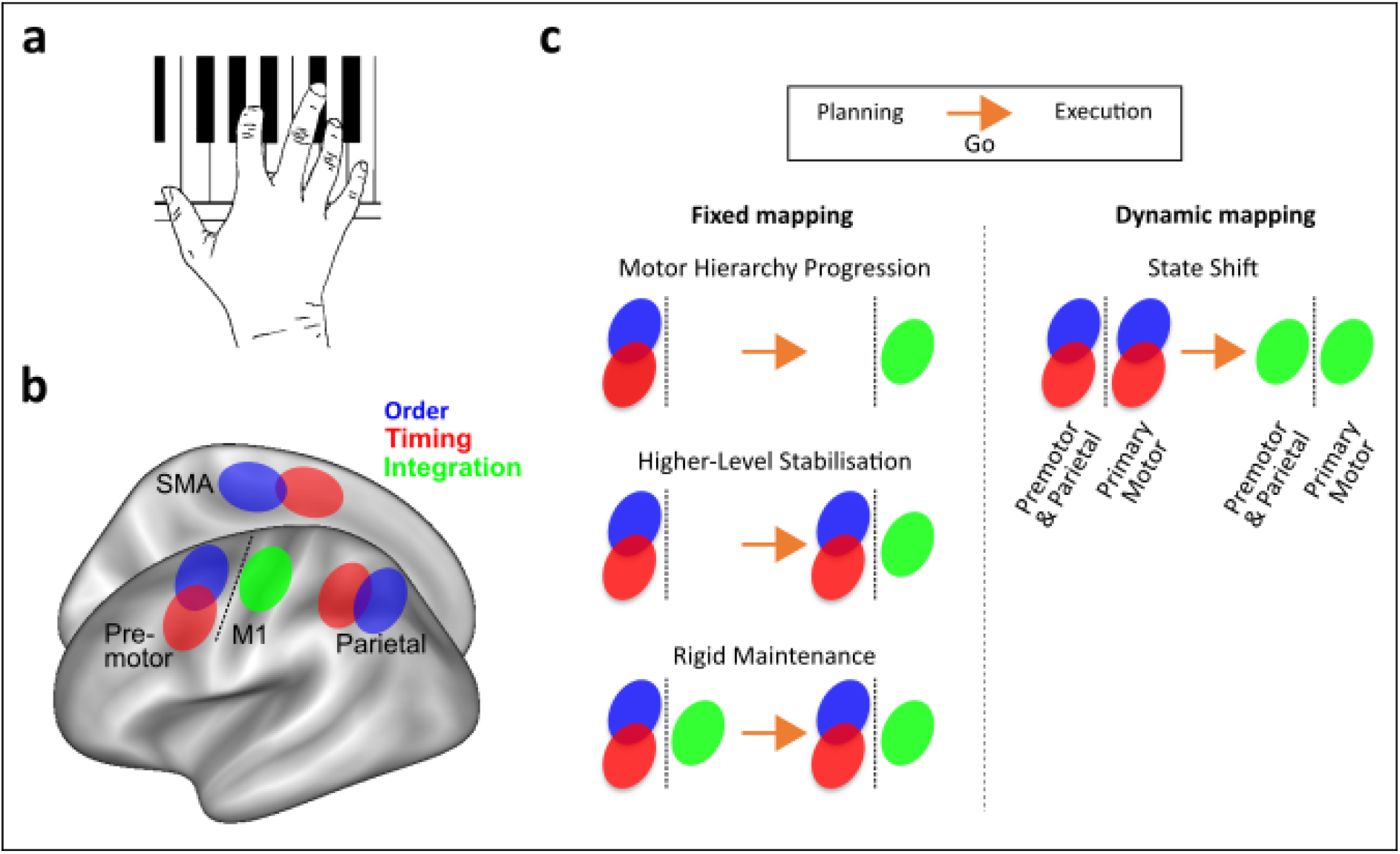
Theoretical framework and hypotheses. (a) Skilled sequence production, e.g. when playing a melody on a piano, is characterised by producing movements with a specific order and timing and combining them flexibly trial-by-trial. (b) Previous findings localised independent patterns of order and timing to premotor, supplementary motor, and parietal regions, while their integration was found in the primary motor cortex (M1) ^12^. (c) How does this mapping evolve across planning and execution? “Fixed mapping” hypotheses state that premotor and parietal regions outside of M1 control order and timing as independent motor sequence features, and M1 itself controls the non-linear integration of the two during planning and/or production. In contrast, the “Dynamic mapping” hypothesis states that there is a state shift within regions from independent feature control during planning to integration during execution (OSF preregistration: https://doi.org/10.17605/OSF.IO/G64HV).

Previous behavioural ^4–6^, computational ^7,8^, neurophysiological ^9–11^ and neuroimaging findings ^12–14^ have established that movement order is controlled independently of timing, and vice versa, whenever motor sequences incorporated temporally discrete sub-goals. This includes sequences that are extensively trained and performed from memory without external guidance, characteristic of motor sequence execution in the real world. The integration of movement timing and order has been studied in the context of execution ^6,12,15–17^, but we currently do not know whether movement order and timing integration takes place prior to movement onset, and which brain regions hold this information about the upcoming sequence.

Neural and haemodynamic activity patterns in contralateral primary motor and sensorimotor, premotor, and parietal cortices show informational tuning to trained motor sequences ^12,18–25^. Specifically, activity patterns in motor-related regions outside the primary motor cortex - the bilateral premotor, supplementary motor and parietal areas - contain high-level information, e.g., about sequence chunks and positional rank in the sequence ^18,26,27^ and spatial, rather than body-centred coordinates ^22^. Further, activity patterns in these regions can generalise across different pairings of movement order and timing ^12^(Figure 1b). In contrast, activity patterns in contralateral primary motor (M1) and sensorimotor (S1) cortices contralateral to the movements are associated with the planning and execution of single movements in a sequence ^11,20,21,28^ and are largely effector-specific ^22^. Further, activity in M1 has been shown to hold information about unique sequence order and timing combinations, suggesting a lower-level integrated representation of the sequences ^12^ (Figure 1b), but may also be reflecting patterns that are driven by differences in the identity of the first movement in a sequence ^20,26^.

Despite the progress made, we lack knowledge of when and where motor-related cortical areas integrate the order of movements with their timing for successful sequence production. One possibility is that these regions show a fixed mapping to high-level independent (premotor, parietal) or lower-level integrated (primary motor and sensorimotor) features of sequences, respectively ^12^. These may be activated simultaneously or sequentially depending on the peri-movement phase, but their informational content would remain stable (Figure 1c, “Fixed mapping”). Alternatively, the tuning of these regions to high and low-level features of sequences may change dynamically depending on trial phase, such that the same regions parse sequence order and timing during planning but integrate these sequence features during execution (Figure 1c, “Dynamic mapping”).

Here, we trained participants to produce five-element finger press sequences comprised of two finger orders and two temporal interval orders (timings) from memory in a delayed production paradigm. To disentangle planning from execution using fMRI, activity was extracted from ‘No-Go’ and ‘Go’ trials, respectively. We utilised multivariate pattern analysis to distinguish between independent (transferrable) and integrated (unique) BOLD patterns related to the planned and executed sequence order and timing. Our results provide strong evidence for the integration of sequence order and timing during sequence execution only, but not during planning. Further, they support the idea that contralateral premotor to parietal regions are not fixed in their informational content but update their tuning dynamically.

## Results

### Discrete sequence production from memory

Participants were trained to produce four finger-press sequences from memory with the right hand on a force transducer keyboard (Figure 2a). Training consisted of a three-staged transition across two days from trials which visually guided sequence production, towards trials which required sequence production entirely from memory (Figure 2b). During functional MRI scans taking place on the third day, participants were required to produce movement sequences from memory only (see supplementary table S1 for trial distribution). Sequences were cued 1000-2500ms before the Go cue by a Sequence cue (abstract fractal image) to prompt the planning of the respective sequence without movement (Figure 2c). To isolate fMRI responses to movement planning without contamination from execution patterns, in addition to ‘Go’ trials, ‘No-Go’ trials were implemented which consisted only of the Sequence cue but did not contain a Go cue (Figure 2d). ‘No-Go’ trials made up 20% of trials during training, and 50% of trials during the fMRI session (see Methods). The target sequences were unique combinations of two finger orders consisting of five presses matched in finger press occurrence and two target relative inter-press-interval (IPI) orders involving four IPIs matched in target duration (Figure 2e). The finger orders were generated pseudo-randomly for each participant, but each sequence started with the same finger press within each participant to avoid first-finger identity driving the sequence decoding during the preparatory period ^20^. Timing 1 and Timing 2 were the same across participants.

**Figure 2.**
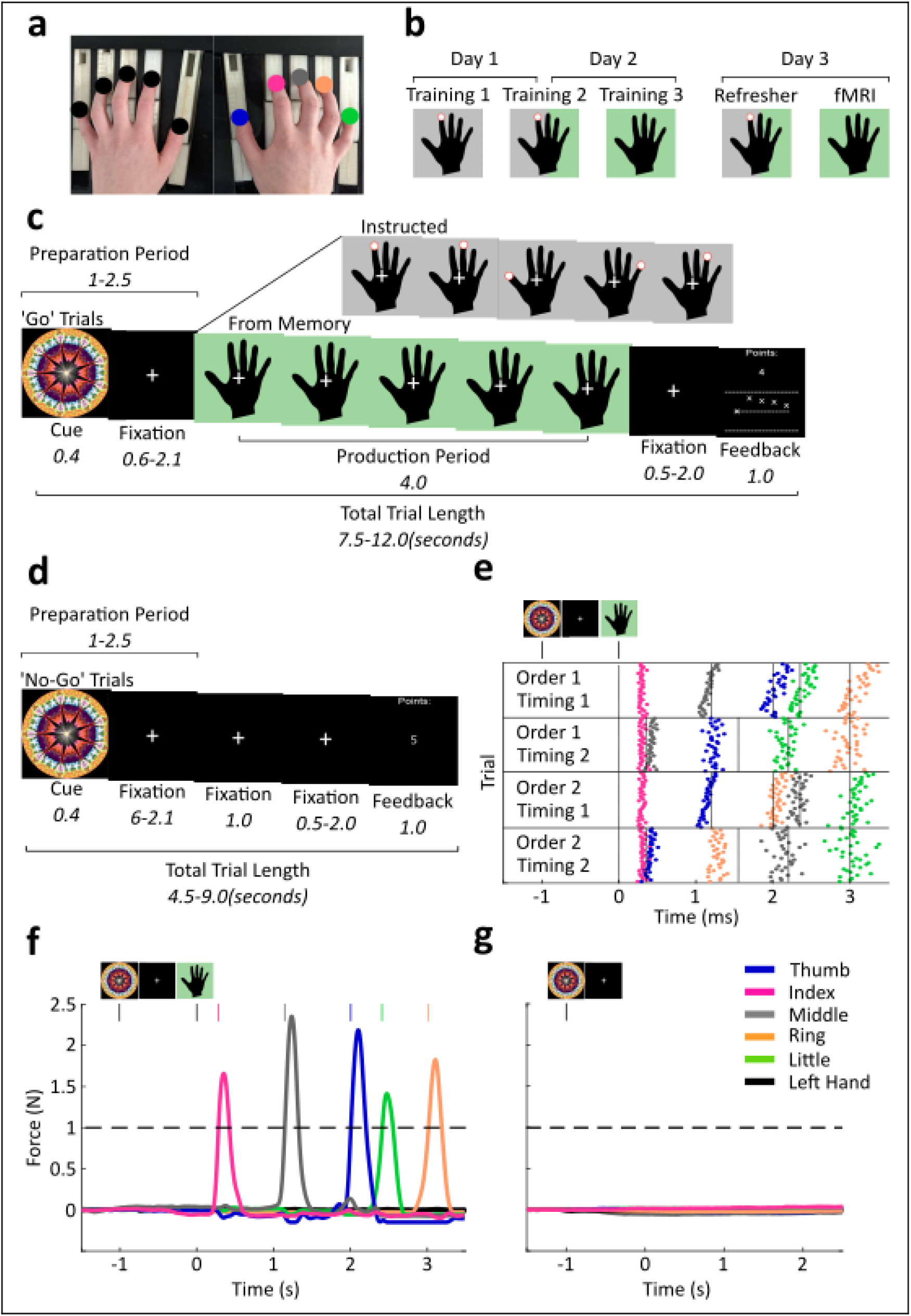
Experimental and trial designs. (a) Participants produced finger presses on a ten-finger force transducer keyboard. The hands were visually occluded from the participants’ view by a panel during training and when lying in a supine position during the fMRI session. Target fingers on the right hand are indicated by different colours that also correspond to the legend in later panels. Fingers on the left (inactive) hand are marked as black. (b) Trial type proportions on each experimental day progressed from 100% instructed (Training 1) to 50/50% mixed (Training 2) to 100% from memory (Training 3) trials during the last stage of training and during fMRI. Black hands with a grey background and a red finger cue indicate visually instructed trials. Black hands with a green background represent trials with sequence production from memory. (c) ‘Go’ trials from memory consisted of a Sequence cue, followed by a fixation cross and a Go cue instructing a production period. The occurrence of the Go cue was the onset of the respective hand stimulus. The trial ended with a feedback screen which indicated finger and temporal accuracy relative to a target sequence. Instructed trials, shown as an insert at the top of the image, followed the same trial structure as from memory trials, but displayed visual finger cues to aid production. (d) ‘No-Go’ trials consisted of a Sequence cue, followed by a fixation cross without a Go cue, and feedback screen. (e) A raster plot shows all button press timings in correct trials produced from memory across the entire fMRI session in one representative participant. Horizontal lines separate the different sequences that followed a two finger order by two timing design (see Methods for details). Vertical dotted lines indicate target press timings. Each coloured dot represents a different effector, see corresponding legend. (f) Example force traces from 10 channels corresponding to the fingers on the right (coloured) and left hands (black) in one representative ‘Go’ trial during fMRI. The horizontal dashed line represents the finger press threshold, and coloured vertical lines represent the time point at which a press was detected from the respective finger. (g) Example force traces, as in f, from one representative ‘No-Go’ trial.

The keyboard recorded isometric force trajectories from fingers of both the active right and the passive left hand concurrently during preparation and production (Figure 2f, Figure 2g). Points were awarded trial-by-trial only if participants did not exceed a force threshold above the baseline period during preparation and ‘No-Go’ trials. In ‘Go’ trials points were calculated based on initiation time after the Go cue, finger press accuracy, and timing accuracy. ‘No-Go’ trials were rewarded when no responses were made above threshold (see Methods). To ensure that participants were not pre-pressing the keys below the force threshold, we checked offline if exerted force of the right hand increased significantly above the baseline level. In ‘No-Go’ trials we checked for force increase from the Sequence cue onset to the last possible moment a Go cue could appear if it were a ‘Go’ trial, to represent the preparatory period. Participants did not increase force during ‘No-Go’ trials, and instead showed a small but significant force reduction (*M* = 0.154N, *SD* = 0.09) relative to baseline (*M* = 0.162 N, *SD* = 0.09; *t*(23) = 3.35, *p* = .003). A similar small decrease, rather than an increase, was found in the preparation phase of ‘Go’ trials (*M* = 0.163 N, *SD* = 0.09) relative to baseline (*M* = 0.164 N, *SD* = 0.09; *t*(23) = 2.44, *p* = .023), suggesting that this force decrease associated with planning was not specific to ‘No Go’ trials. Importantly the data shows that participants did not engage in any subthreshold pre-pressing or rehearsal of the sequence during sequence preparation.

All included participants reached the criteria of producing different timing structures (Timing 1 and 2) across finger orders (Order 1 and 2; Figure 3a shows group-level data, ANOVA results in Methods). We did not expect any main effect or interactions with finger order to be significant at the group level, due to the randomisation of trained finger orders across participants (see Methods). However, we assessed the main effects of order and timing, and their interaction at the individual level. Here 18 out of the 24 participants showed a significant order by interval position interaction, and 10 showed a significant three-way interaction between timing, order, and interval position. The presence of these idiosyncratic press timing patterns at the individual level suggests the integration of sequence order and timing features. Crucially, sequence timing error showed no difference between timing structures suggesting that there were no systematic differences in difficulty for Timing 1 and Timing 2 at the group level (*F*(1,23) = 0.07, *p* = .792, *ηp^2^* = .003; Figure 3b). At the individual level, 10 participants showed a significant main effect of order, 15 showed a significant main effect of timing, and four showed a significant interaction between order and timing, again suggesting an integration of the two sequence features.

**Figure 3.**
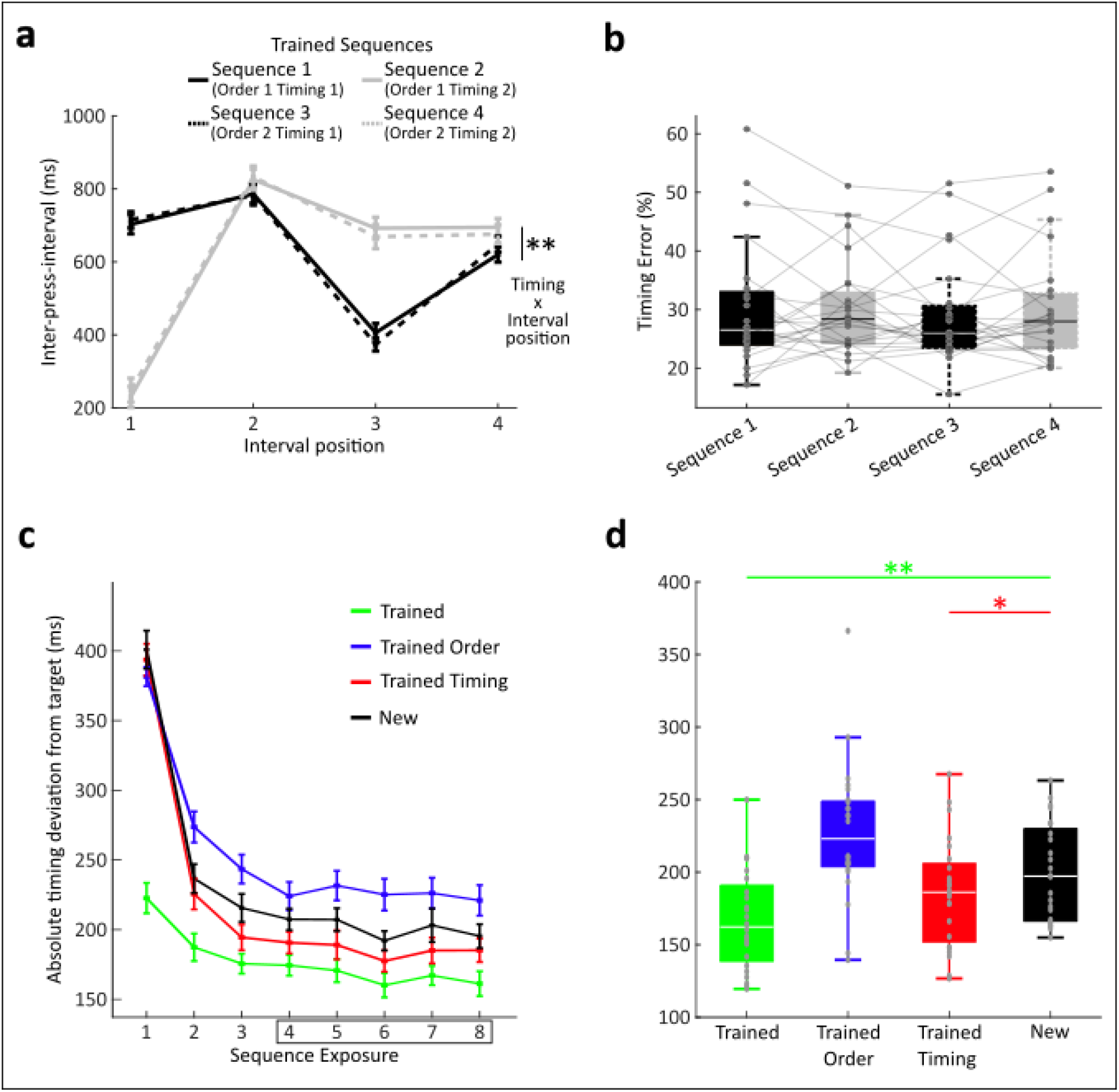
Sequence timing and feature transfer. (a) Inter-press-interval (IPI) structure of the four trained sequences during the fMRI session (day 3). A significant interaction between sequence timing and interval position shows distinct IPI sequences between Timing 1 and Timing 2 across finger order sequences Order 1 and 2. (b) Timing error (normalised to the target interval durations, see Methods) during the fMRI session did not differ between sequences. (c) Behavioural transfer results from the synchronisation task obtained from instructed ‘Go’ cue trials following the last training stage on day 2. Absolute deviation from target timing are shown across sequence repetitions for trained sequences (green), sequences with trained finger orders, but unfamiliar timing (blue), sequences with trained timing, but unfamiliar order (red), and new sequences with both unfamiliar finger order and timing (black). (d) Absolute deviation from target timing, as in a, extracted from the fourth to the last sequence repetition as in previous work (e.g., Kornysheva et al. 2019). Significance of t-tests to identify performance benefits compared to new sequences is shown by coloured asterisks and horizontal lines. Note that the trained order condition showed a significant increase in synchronisation error (*p* = .002, two-tailed t-test), suggesting interference rather than benefits related to sequence feature transfer. ** *p* < 0.01; * *p* < 0.05, one-sided t-test.

To test for independent transfer of finger order and timing to new combinations before and after training, participants performed a synchronisation task where we assessed their synchronization error (absolute deviation from target timing) to a visually cued sequence ^14^.The trials in each condition were presented in a blocked manner with 8 repetitions to assess short-term learning gains related to trained finger order and timing (Figure 3c). Since the transfer of trained sequence timing to a new finger order only takes place after three exposures, synchronisation performance was only assessed from the fourth sequence exposure onwards consistent with previously reported analyses ^6,12,14^. During the post-training phase, we compared each condition (trained, order transfer, timing transfer) to completely new sequences (*M* = 196.15ms, *SD* = 34.00) in a one-tailed paired sample t-test, with trained sequences (*M* = 160.43ms, *SD* = 33.09) showing a significant performance advantage, as predicted (*t*(23) = 6.34, *p* > .001), as did timing transfer sequences (*M* = 182.10, *SD* = 39.72; *t*(23) = 2.09, *p* = .024), replicating previous findings ^6,12,14^ (Figure 3d). In contrast to earlier reports, order transfer sequences (*M* = 223.87, *SD* = 48.30) showed a significantly worse performance (*t*(23) = 3.52, *p* = .002, two-tailed test). Whilst knowledge of both features of a sequence combined, or just its timing, facilitated task performance, knowledge of sequence order hindered future learning of novel sequence acquisition when paired with a new timing structure. This implies that the current participants acquired a stronger independent representation of timing than of finger order which was integrated with a particular timing structure during production.

### Activity increases during preparation and production

Percent signal change of preparation and production relative to rest were calculated to identify areas with increased activity during these two respective phases of the task. While preparatory activity was modelled in the GLM for both ‘Go’ and ‘No-Go’ trials, preparatory activity for % signal change and multivariate pattern analyses was solely sampled from No-Go trials to disentangle the BOLD activity in these two phases whilst keeping a fast event-related design. We then calculated the percent signal change across the cortex (Figure 4a) and extracted values along two cross-sections of the cortical surface on the contralateral (left) side to the motor effector ^12^ (Figure 4b). These cross-sections extended from anterior to posterior and ventral to dorsal, across premotor to parietal and premotor to supplementary motor regions respectively, because our hypotheses (osf.io/g64hv) on the imaging results during sequence preparation and production were put forward for contralateral premotor, primary motor, and parietal regions, which we expected to be tuned to sequence information based on previous studies ^12,18–25^. Whole-brain results are presented in the Supplementary Materials. We carried out one-tailed non-parametric permutation tests along these cross-sections to identify significant clusters where activity increased above baseline ^29^. During preparation, a very small, but significant, activity increase was found within ventral premotor cortex (PMv) (*p* = .002), and a marginally significant increase found within dorsal premotor cortex (PMd) (*p* = .050) (Figure 4b). During production, significant activity increases were found across the majority of contralateral motor-related regions, with one large cluster across PMd, M1, sensorimotor cortex (S1), anterior superior parietal cortex (SPCa), and posterior superior parietal cortex (SPCp) (*p* < .001), and another cluster which spanned the cross-section from PMv to PMd (*p* < .001). The cross-section overlapping with anterior supplementary motor area (SMA) did not show a significant activity increase from rest during production. However, note that the section of the SMA directly posterior to the cross-section did show a significant activity increase (see supplementary Figure S1a and Table S2 for whole-brain contrast cluster analysis).

**Figure 4.**
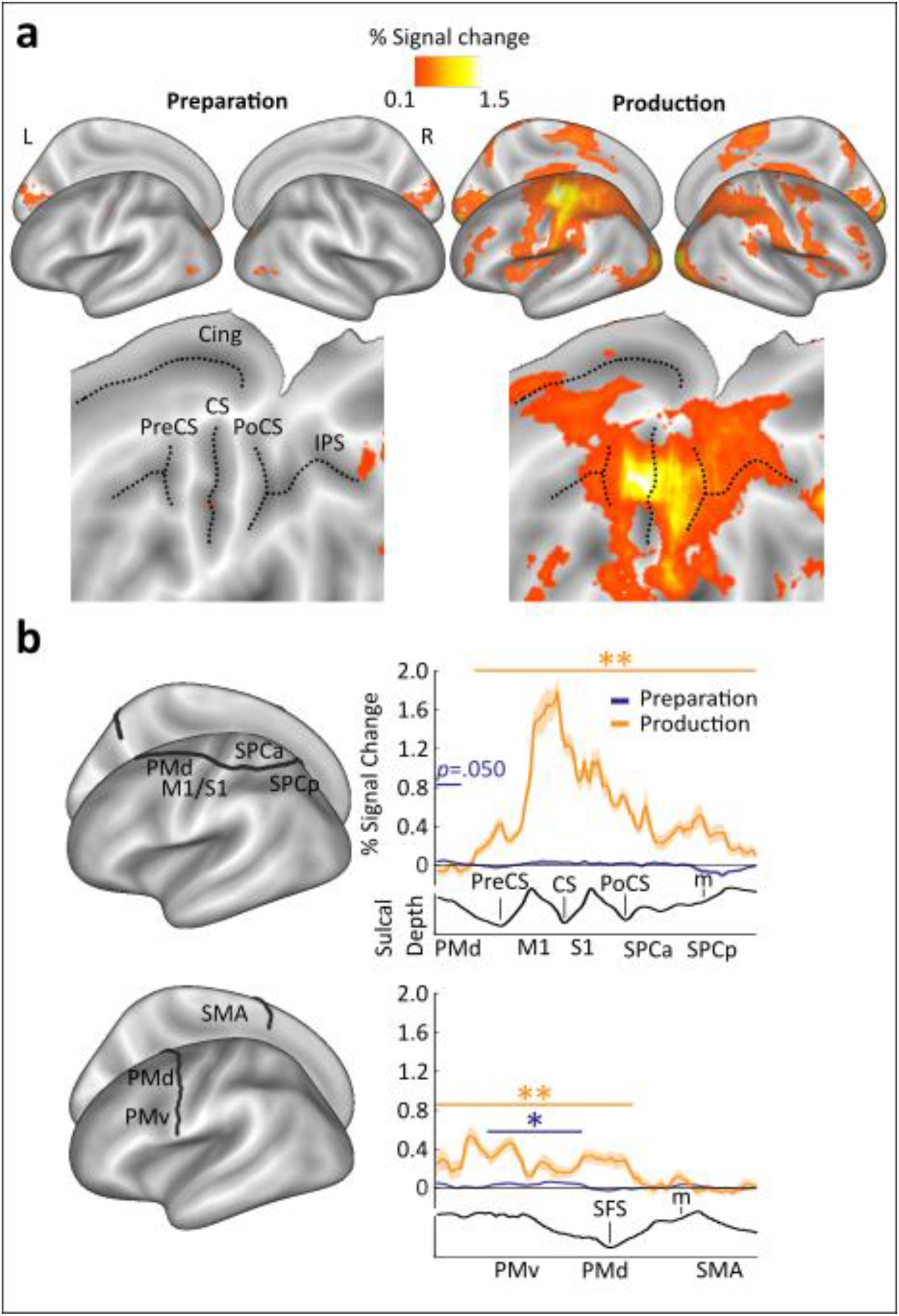
Percent signal change during preparation and production. (a) Inflated surface maps are shown in upper panels and flat maps in lower panels, displaying mean % signal change during preparation (left panels) and production relative to rest (right panels), respectively. (b) Mean % signal change relative to rest for both preparation (blue lines) and production (orange lines), plotted on cross-sections running from rostral premotor cortex, through the hand area, to the occipito-parietal junction (upper panel) and on a profile running from the ventral, through the dorsal premotor cortex, to the SMA (BA6; lower panel). Clusters with significant increases above baseline are denoted by the coloured horizontal lines and asterisks, calculated using one-tailed non-parametric permutation tests. Cing, cingulate; CS, central sulcus; IPS, intraparietal sulcus; m, medial wall; M1, primary motor cortex; OPJ, occipito-parietal junction; PMd, dorsal premotor cortex; PMv, ventral premotor cortex; PoCS, post-central sulcus; PreCS, pre-central sulcus; pre-SMA, pre-supplementary motor area; S1, sensorimotor cortex; SFS, superior frontal sulcus; SMA, supplementary motor area; SPCa, anterior superior parietal cortex; SPCp, posterior superior parietal cortex. ** *p* < 0.01; * *p* < 0.05, one-sided t-test.

### Multi-variate pattern analysis (MVPA)

We used MVPA to identify cortical areas that showed systematic changes in BOLD activity patterns between sequences during preparation and production. Using a whole brain searchlight of 160 voxels ^30^, we trained a linear discriminant analysis (LDA) classifier to distinguish between sequences in a one-run-out cross-validation method – an approach that has been validated with pattern simulations in a previous study ^12^. Specifically, we looked for regional activity patterns that either transferred across or were unique for specific combinations of order and timing. The order classifier was used to decode between sequences with different finger orders, regardless of their pairing with a timing feature, whereas the timing classifier was trained to decode between sequences with different finger timings regardless of their pairing with a specific finger order. These two classifiers allowed the identification of regions which contained above chance decoding of sequence order and timing independently of the other sequence feature, respectively (Figure 5a and Supplementary Figure S2 for t-maps). The integrated classifier decoded residual patterns after subtracting averaged sequence order and timing related patterns for each run separately, in order to detect regions which hold information on sequence identity that is not driven by a simple summation of order and timing information (see Methods).

**Figure 5.**
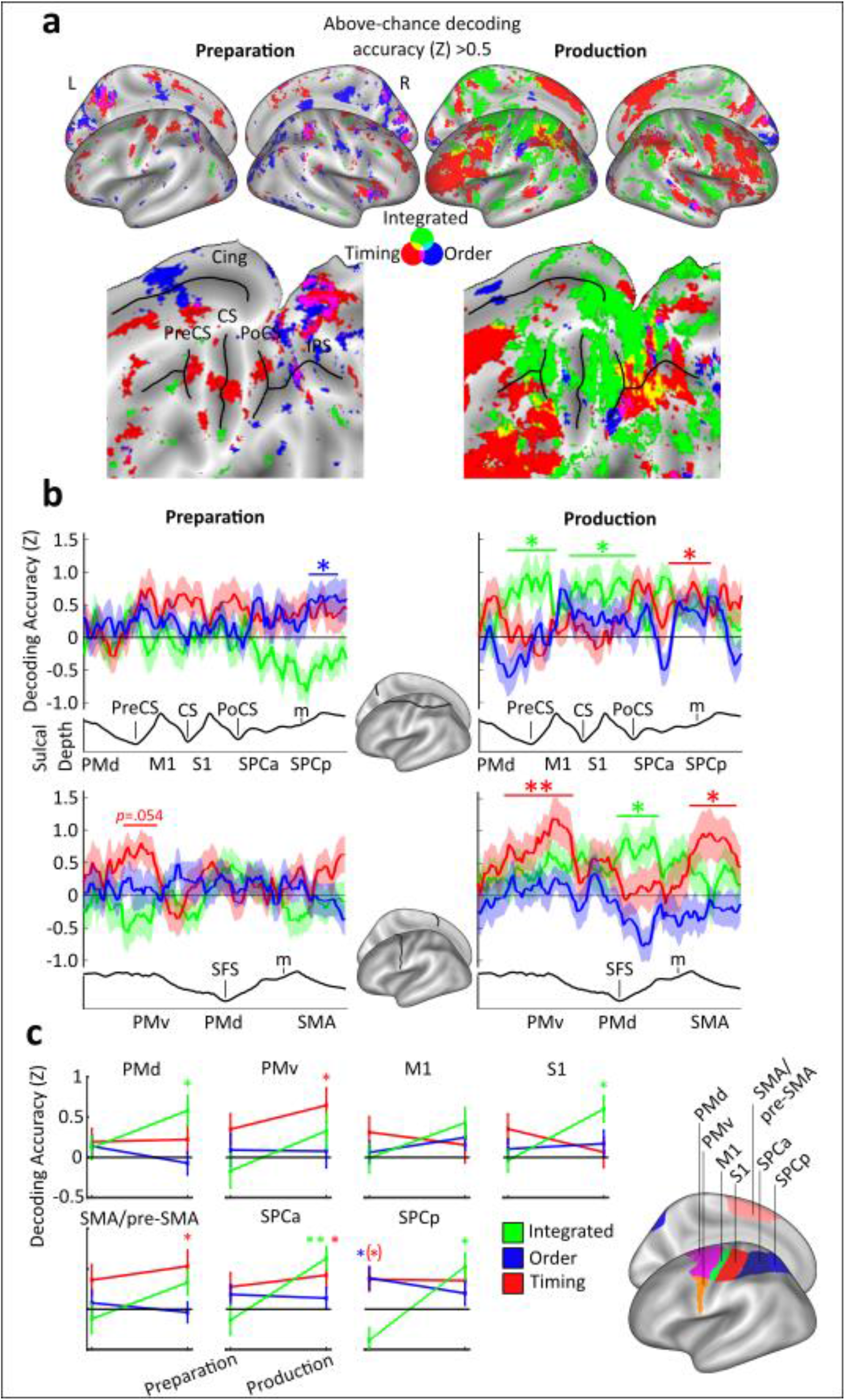
Multivariate pattern classification results. (a) Inflated surface (upper panels) and flat maps (lower panels), showing mean decoding z-accuracy values above chance for finger order (blue), timing (red), and integrated sequence patterns (green). (b) Mean decoding z-accuracy values for each classifier along the cross-sections explained in Figure 4b. Coloured asterisks denote the respective significant above-chance clusters for each classifier during preparation and production. (c) Searchlight z-accuracy values were extracted using pre-determined ROIs shown in the left panel. Decoding above is indicated by the coloured asterisks. The asterisk in parentheses represents a marginally significant result, *p* = .060. ** *p* < 0.01; * *p* < 0.05, one-sided t-test.

To reveal the continuous profile of feature decoding along contralateral motor regions on the cortical surface, we employed the same permutation test approach ^29^ as in the % signal change analysis for each of the three classifiers, for preparation and production, separately (Figure 5b). During preparation, a significant cluster was found for finger order within SPCp (*p* = .040), and a marginally significant cluster for timing decoding was identified within PMv (*p* = .054). During production, above chance decoding was shown for the integrated classifier within PMd in two clusters (*p* = .002, *p* = .044, on anterior to posterior and ventral to dorsal cross-sections, respectively) and S1, which extended into M1 and SPCa (*p* = .007). Above chance decoding of timing was found within SPCa (*p* = 0.016), PMv (*p* < .001), and SMA (*p* = .045).

Next, we examined how well sequence features could be decoded from pre-registered ROIs during preparation and production. These regions covered premotor to superior parietal areas: PMd, PMv, M1, S1, SMA/pre-SMA, SPCa, and SPCp. First, to identify above chance decoding of sequence information in these areas, one-sample t-tests were performed on the z-values extracted from each of the pre-defined ROIs during both preparation and production for timing, order, and integrated classifiers (Figure 5c). These t-tests were Bonferroni corrected six times, to account for phase (2) by classifier (3) within each pre-defined ROI. During preparation, the only region that reached significance above chance was found in SPCp for sequence order decoding (*t*(23) = 2.74, *p* = .036), with marginally significant decoding also in SPCp for sequence timing (*t*(23) = 2.51, *p* = .060). During production, classification increased above chance for sequence timing in SMA/pre-SMA (*t*(23) = 2.71, *p* = .036), PMv (*t*(23) = 3.00, *p* = .018), and SPCa (*t*(23) = 2.67, *p* = .042). Further, classification increased above chance for order-timing integration in S1 (*t*(23) = 3.69, *p* = .003), PMd (*t*(23) = 3.06, *p* = .018), SPCa (*t*(23) = 4.36, *p* <.001), and SPCp (*t*(23) = 3.20, *p* = .012).

Finally, we set out to test our main hypotheses (osf.io/g64hv) regarding an interaction between peri-movement phase (preparation, production), classifier (timing, order, integrated) and region (PMd, PMv, M1, S1, SMA/pre-SMA, SPCa, SPCp). A repeated-measures ANOVA revealed a significant main effect of phase (*F*(1,23) = 9.49, *p* = .005, *ηp^2^* = .292), substantiating a general increase of decoding accuracy across regions and classifiers during production. The main effect of region was not significant (*F*(3.84,88.42) = 0.45, *p* = .763, *ηp^2^* = .019, Greenhouse-Geisser corrected), suggesting that all the contralateral cortical ROIs had a comparable contribution to sequence decoding across trial phases. Importantly, we found a significant phase by classifier interaction (*F*(2,46) = 10.34, *p* = .044, *ηp^2^* = .127), which was driven by an overall increase in the integrated classifier accuracy from preparation (*M* = −0.10, *SE* = 0.13) to production (*M* = 0.49, *SE* = 0.11) (*p* = .003, *95% CI* [.217, .971], Bonferroni corrected). Finally, there was no significant interaction of phase by region (*F*(3.20,73.50) = 0.79, *p* = .512, *ηp^2^* = .033, Greenhouse-Geisser corrected), or phase with classifier by region (*F*(5.40,124.18) = 1.63, *p* = .151, *ηp^2^* = .066, Greenhouse-Geisser corrected). In sum, this supports the hypothesis that tuning of these regions to high and low-level features of sequences changes dynamically depending on trial phase, rather than region, with a state shift towards sequence feature integration after movement initiation across multiple regions.

## Discussion

Activity in the premotor, primary motor and parietal cortices has been associated with motor sequence control, from their hierarchical organisation ^11,17,22,26,27,31,32^ to the control of spatial and temporal features of sequences ^9,12,22,33–35^. Yet how sequence-related computations in these cortical regions unfold across planning and execution phases remains uncertain. Do these cortical areas retain a fixed tuning to sequence features and their integration throughout planning and execution? Or do they switch their content dynamically depending on whether they occur before or after motor initiation? Here, we examined how motor cortical areas integrate informational content on sequence features – the order of finger movement sequences and their timing – across the planning and execution phases. Sequence decoding from local cortical activity patterns revealed that high-level features of sequence organisation remain separate during movement planning and are integrated holistically into unique patterns upon movement initiation in premotor and parietal areas.

### Cortical patterns switch their tuning from planning to execution

Our results demonstrate a generalised dependency of cortical representations on peri-movement phase, with a global shift across regions towards feature integration at the transition from sequence planning to execution. This indicates that cortical motor-related areas do not rigidly map onto higher-level versus lower-level representations of sequential organisation, respectively, as assumed by earlier studies that focussed on activity patterns during sequence execution alone ^12,26,36^. Instead, pattern activity tuning in these regions changes dynamically upon motor initiation. Such a state switch from higher-level independent to lower-level integrated control echoes previous findings for the primary motor and dorsal premotor cortices in the context of single movements. These show that preparatory neural population activity occupies a different state space (output-null) from the production activity to prevent readout from downstream areas during planning ^11,37,38^. Here, cortical motor planning patterns are not simply subthreshold versions of execution activity patterns controlled by inhibitory gating within the cortex or downstream as suggested previously ^39,40^, but a qualitatively different activity pattern of the neural population. Our results support the notion of a distinct functional tuning during motor planning across regions on the premotor to parietal axis in the context of sequential movements.

### Lack of sequence feature integration prior to motor initiation

As participants trained to perform the four finger sequences over two days and entirely from memory, one may expect that this level of practice would result in their retrieval as one integrated spatio-temporal synergy ^41^. However, we found that information about motor sequence order and timing of the upcoming sequence was reinstated independently and integrated after motor initiation only. This recompilation of sequence features occurred on a trial-by-trial basis. One possibility is that the observed lack of integration during preparation may be due to requirements of the two-by-two task design, which may encourage participants to hold onto higher level representations of order and timing. However, in previous tasks where only one combination of timing and order was trained, independent transfer of known order and timing to new combinations was still found, showing that the separation is not dependent on the task structure, but occurs more generally ^5,6^.

Although previous work has shown that planning-related activity in motor areas is predictive of movement features such as speed, force, and trajectory of the upcoming movement ^42–44^, these may be regarded as part of planning a holistic motor synergy ^45–47^. In contrast, for sequence learning there is now ample evidence that higher-level sequence features such as movement order and timing are encoded independently ^5,7,12,13^ and remain separate during planning ^14,48^, despite training across multiple days and production from memory. This is in line with recent evidence in macaque motor cortex suggesting that the neural generation of sequence elements with a discrete timing goal – separated by a delay or in rapid succession – remains independent, despite long-term training and fusion at the muscular level ^11^.

Our results show a clear phase-related state shift within regions for discrete movement sequences features, yet how this integration occurs in more continuous overlapping movement sequences is unclear. Rather than a dedicated timing system as is observed with discrete movements ^5,12,13,16^, continuously overlapping movements have been shown to employ a more state-dependent timing system ^49–52^ meaning that they are likely to employ differing neural mechanisms. Further research should investigate whether sequential movements with overlapping, continuous trajectories that lack separate timing goals show the same state shift in informational tuning observed in the current findings.

What triggers sequence feature integration trial-by-trial? We propose that contralateral motor-related cortical regions activate movement order and timing plans separately until a sensory stimulus like the Go cue triggers the binding of the corresponding neural patterns. This binding may occur through subcortical, e.g. thalamic input triggering an appropriate state for motor execution of specific combination of features ^53,54^. In sum, delaying the binding of sequence features to the production phase and maintaining higher level separation may allow the system to retain maximum flexibility trial-by-trial, should task demands change.

### Independent patterns for sequence timing are reinstated during execution

We found a stark asymmetry with regard to sequence order and timing during the sequence production phase. In contrast to the independent patterns for finger order, the activity patterns tuned to sequence timing became more prominent or emerged during production in the PMv and SMA/pre-SMA, as well as the superior parietal cortices. Thus, cortical patterns for sequence timing are reinstated alongside the emergence of sequence-specific integrated patterns. This asymmetry was also observed at the behavioural level in the transfer task. Here, trained timing could be quickly recombined with a new finger sequence producing significant advantages to a sequence that had both new timing and new order in line with previous work ^6,12,14–16^. In contrast, producing the same finger order with a new timing was associated with a significantly poorer performance. Thus, in contrast to a previous study utilising the same task paradigm and well-trained sequences produced from memory ^14^, here, participants were unable to separate the trained order from their timing. This interference effect directly parallels the prominence of integrated and the lack of independent finger order tuning during motor production.

### M1 lacks information about sequences despite a large activity increase during execution

Our results show a lack of sequence feature separation or integration in contralateral M1 during preparation and only limited integration above chance during production extending out from the greater peak in S1. This occurs despite a large activity increase in M1 during production. While this contrasts with several previous neuroimaging studies ^12,19,26,55,56^, recent findings show that information held within M1 is not related to sequence control. Activity patterns do not change with sequence learning ^21,57^ and reflect the processing of individual movements, particularly, the first press of a sequence ^20^. In our study, the first finger press was matched across sequences within participants and the GLM of the BOLD patterns predicted the response function related to this matched first press only, reducing the possible influence of subsequent presses. So far, there has been no experimental evidence that sequential movements are neurally fused in M1 into holistic representations: Constituent movements remain individuated in M1 regardless of sequential context ^11,27^. Therefore, controlling for the first finger press might explain why we see no prominent sequence feature integration within M1, as observed previously ^12^.

### Extending the previous motor planning framework to sequential actions

The current framework for single movement motor planning proposes that the motor system enters a preparatory state that is distinct from movement execution and, when movement is cued, dynamically and passively evolves across a time period in order to produce movements ^37,47,58^. Recent findings also suggest a distinction between the selection of motor goals and true motor planning, which formulate ‘what’ movements to execute and ‘how’ to execute them, respectively ^59,60^, converging with the idea of hierarchical motor control ^1,26^. In the context of skilled sequence control, we propose that sequence order and timing features are specified during planning as ‘what’ elements representing higher-level control, and integrated during execution as ‘how’ elements, representing lower-level implementation motor control. Crucially, our results suggest that individual regions can undergo a state shift from ‘what’ to ‘how’ control depending on the peri-movement phase. Future electrophysiological research should address whether the same neuronal populations are involved in both types of control and determine the neural origin and exact time point that triggers this cortical state shift.

## Materials and Methods

### Participants

24 neurologically healthy participants - 14 females and 10 males (*M* = 21.00 years, *SD* = 1.64) - met all behavioural and imaging requirements after completing the three-day experiment. 23 Participants were right-handed with a mean Edinburgh Handedness Inventory (EHI;https://www.brainmapping.org/shared/Edinburgh.php; adapted from Oldfield (1971)) score of 75.22 (*SD* = 20.97, Range: 25-100), one was left-handed with an EHI score of −70. Data were collected from an additional 17 participants but were excluded. One participant was excluded due to unforeseen technical difficulties with the apparatus and one participant was excluded due to a corrupted functional scan. 15 further participants did not reach target performance during training and so were also excluded following training. Target performance consisted of an error rate below 20% (*M* = 6.54%, *SD* = 6.03, for the group) and distinct sequence timing structures. Distinct sequence timing was fulfilled if there was a significant interval position by sequence timing interaction on IPI duration in each participant during the fMRI session (Group *F*(1.78,40.77) = 73.76, *p* <.001, *ηp^2^* = .762, Greenhouse-Geisser corrected, repeated measures ANOVA). Participants were recruited either through social media and given monetary payment, or through a participation panel at Bangor University and awarded module credits for their participation. Participants with professional musical qualifications were excluded from recruitment. All participants provided informed consent, including consent to data analysis and publication, through an online questionnaire hosted by Qualtrics (Qualtrics, Provo, UT). This experiment and its procedures were approved by the Bangor University School of Psychology Ethics Committee (Ethics approval number 2019-16478).

### Apparatus

Force data from fingers of both the right and left hands were recorded at a sample rate of 1000hz using two custom-built force transducer keyboards (10 channels). Each key had a groove within which the respective fingertip was positioned. A force transducer (Honeywell FS Series, with a range of up to 15 N) was located under each groove and recorded the respective finger force without crosstalk between channels. Force data acquisition occurred in each trial from 500ms before sequence cue onset to the end of the production period in production trials, and the end of the false production period in No-Go trials. The keys could be adjusted in position by sliding them up and down individually along the keyboard plane to achieve the most comfortable position for the hand and wrist when seated during training or in supine position in the MRI scanner, respectively. Once adjusted the position of the keys was fixed. Traces from the right hand were baseline-corrected by the first 500ms of acquisition (500ms before the sequence cue) and smoothed to a Gaussian window of 100ms, trial-by-trial. Button presses were defined as the point at which forces above baseline exceeded a fixed threshold (2.5 N for the first 8 participants and 1 N for the subsequent 16 out of 24 participants). Press timings were identified by the timestamp provided by National Instruments Data Acquisition Software (National Instruments, Austin, TX) associated with the data point at which the respective threshold was exceeded.

During behavioural training sessions, participants were seated at a wooden table approximately 75cm away from a 19-inch LCD LG Flatron L1953HR, at a resolution of 1280 x 1024, at a refresh rate of 60Hz. Their hands were occluded by a horizontally positioned panel on posts around the force boxes. During fMRI sessions, stimuli were presented on an MR Safe BOLDScreen 24”, at a resolution of 1920×1200 and a refresh rate of 60Hz. Participants laid supine on the scanner bed and the two force transducers were positioned on a plastic support board resting on their bent upper legs to enable comfortable and stable positioning of the hands.

### Behavioural task

Participants were trained to produce four five-finger sequences with defined inter-press-intervals (IPI) from memory in a delayed sequence production paradigm. ‘Go’ trials began with a fractal image (Sequence cue) presented for 400 milliseconds (ms) which was associated with a sequence. The mapping between fractal image and each sequence was defined randomly for each participant. Following the Sequence cue, a fixation cross was shown to allow participants to prepare the upcoming sequence; display length of this fixation cross was jittered at durations of 600ms, 1100ms, 1600ms, and 2100ms, pseudorandomised across trials within blocks. A black hand with a green background (Go cue) then appeared for 4000ms to cue sequence production. Succeeding the Go cue, another fixation cross was presented in a jittered fashion at durations of 500ms, 1000ms, 1500ms, and 2000ms. Feedback (see feedback section for more details) was then presented to participants for 1000ms, followed by a jittered inter-trial-interval (ITI) duration of 1000ms, 1500ms, 2000ms, and 2500ms. Visually guided (Instructed) ‘Go’ trials during training were presented in the same fashion albeit featuring a Go cue with a grey background, and a red dot on the tip of each finger on the hand image would move from finger to finger in the target production order and in-pace with the target timing structure. ‘No-Go’ had the same structure to ‘Go’ trials, but no Go cue was shown succeeding the preparatory fixation cross. Instead of the Go cue the fixation cross continued to show for an additional 1000ms. As in ‘Go’ trials, this phase of the trial was followed by a fixation cross, feedback and ITI.

Four target sequences consisted of permutations of two finger orders (Order 1 and 2) and two IPI orders (Timing 1 and 2) matched in finger occurrence and sequence duration. Sequence orders were generated randomly for each participant. All trained sequences began with the same finger press to avoid differences in the first press driving the decoding of sequence identity during preparation ^20^. Ascending and descending press triplets and any identical sequences were excluded. Timing structures were the same across participants, to allow for comparison of timing performance across participants. The two trained timing structures consisted of four target IPI sequences as follows: 1200ms-810ms-350ms-650ms (Timing 1), and 350ms-1200ms-650ms-810ms (Timing 2). To assess if participants maintained the target timing structure despite individual tendencies to lengthen or compress overall sequence length, we calculated timing error for each participant relative to their average total production length. This was calculated offline by normalising target and produced IPIs as a percentage of the participant’s average total sequence length during the session across sequences, then calculating the cumulative percent deviation from target for each IPI, averaged across trials.

Feedback was given to participants trial-by-trial on a points-based scale ranging from 0 to 10. Points were based on initiation reaction time (RT) and temporal deviation from target timing calculated as a percentage of the target interval length. For initiation reaction time, up to five points were awarded for a fast initiation RT as follows: five points for presses within 200ms of the Go cue, four points for presses within 200-360ms, three points for presses within 360-480ms, two points for presses within 480-560ms, one point for presses within 560-600ms, and zero points for presses beyond 600ms. For IPI performance, up to five points were awarded based on deviation from target IPI structure in percent of respective interval to account for the scaling of temporal error with IPI length ^62^. Five points were awarded for average deviations of IPIs from target for each trial which was lower than 10 percent, four points for 10-20 percent, three points for 20-30 percent, two points for 30-40 percent, one point for 40-50 percent, and zero points for above 50 percent. If the executed press order was incorrect, participants were awarded 0 points for the trial. If the executed press order was correct, they were awarded their earned timing points. To discourage premature key presses in the preparation period of ‘Go’ trials and ‘No Go’ trials, 0 points were awarded if participants exceeded a force threshold during preparation above the baseline period. In No-Go trials, five points were awarded if no press was made as instructed. A monetary reward of £10 was offered to the two participants who accumulated the most points across the course of the experiment, to incentivise good performance.

Participants were presented with a feedback screen after each trial showing the number of cumulative points across the whole experiment, as well as feedback on whether they pressed the correct finger at the correct time. A horizontal line was placed in the centre of the screen, with four symbols displayed equidistantly along the line which represented each of the five finger presses. An ‘X’ indicated a correct finger press, and a ‘-’ indicated an incorrect finger press for each sequence position. The vertical position of these symbols above (“too late”) or below (“too early”) the line was proportional to the participant’s timing of the respective press relative to target IPI (in %). Using these cues, participants could adjust their performance online to ensure maximum accuracy of sequence production and prevent a drift in performance from memory following training. During the first two days of training, auditory feedback in the form of successive rising tones corresponding to the number of points (0-10) were played alongside the visual feedback. Auditory feedback was absent during the fMRI session, to prevent any auditory processing driving decoding accuracy.

### Procedure

Training duration was fixed across participants and occurred across the first two days of the experiment over three distinct training stages (see Figure 2 for a visual representation of the training stages). In the first training stage, 80% of all trials were instructed ‘Go’ trials (black hand on grey background, Figure 2c), and the remaining 20% were ‘No-Go’ trials. During the second training stage, 40% of trials were instructed ‘Go’ trials, 40% were from-memory ‘Go’ trials (black hand on green background, Figure 2c), and 20% were ‘No-Go’ trials (Figure 2d). In the third and final stage of training, 80% of trials were from-memory ‘Go’ trials, and 20% were ‘No-Go’ trials. Each stage of training consisted of 240 trials for a total of 720 trials across all three training sections. The third and final day consisted of a short refresher stage of 40 trials, made up of the same proportion of trials as the second stage of training, during which T1 images were collected. Following this refresher stage there was a fMRI stage consisting of 288 trials (48 trials in each block) featuring 50% from-memory ‘Go’ trials and 50%‘No-Go’ trials.

In addition, before and after the last training stage, participants completed a synchronisation task during which participants were asked to synchronize their respective presses to a visual finger cue, as in the first stages of training consisting of four blocks of 32 trials which included trained sequences, sequences with new timings but the same orders (order transfer), sequences with the same timings but new orders (timing transfer), and new sequences. Trial structure was identical to instructed ‘Go’ trials. There were four sequences belonging to each condition and each sequence was shown for eight consecutive exposures (Figure 3c) to assess short-term learning gains. We expected that participants would show significantly more accurate synchronization to visual sequences when they encountered trained sequences as well as sequences with a trained finger order or trained timing compared to untrained control sequences following the completion of training.

### MRI acquisition

Images were obtained on a Philips Ingenia Elition X 3T MRI scanner using a 32-channel head coil. T1 anatomical scans were acquired using a magnetisation-prepared rapid gradient echo sequence (MPRAGE) scan at a 0.937 x 0.937 x 1 resolution, with a field of view of 240 x 240 x 175 (A-P, R-L, F-H), encoded in the anterior-posterior dimension.

T2*-weighted Functional images were collected across six runs of 230 volumes each with a TR of 2 seconds, a TE of 35ms and a flip angle of 90°. The voxel size was 2mm isotropic, at a slice thickness of 2mm, with 60 slices. These were obtained in an interleaved odd-even echo-planar imaging (EPI) acquisition at a multi-band factor of two. Four images were discarded at the beginning of each run to allow the stabilisation of the magnetic field. The central prefrontal cortex, the anterior temporal lobe and ventral parts of the cerebellum were not covered in each participant. Jitters were employed within each trial during preparation periods, post-production fixation crosses, and inter-trial intervals, to vary which part of the trial is sampled by each TR and therefore give us a more accurate estimate of the hemodynamic response function ^63^.

### Pre-processing and first-level analysis

All fMRI pre-processing was completed using SPM12 (Revision 7219) on MATLAB (The MathWorks, Inc., Natick, MA). Slice timing correction was applied using the first slice as a reference to interpolate all other slices to, ensuring analysis occurred on slices which represent the same time point. Realignment and unwarping were carried out using a weighted least-squares method correcting for head movements using a 6-parameter motion algorithm. A mean EPI was produced using SPM’s ‘Imcalc’ function, wherein data acquired across all six runs was combined into a mean EPI image to be co-registered to the anatomical image. Mean EPIs were co-registered to anatomical images using SPM’s ‘coreg’ function and their alignment was checked and adjusted by hand to improve the alignment, if necessary. All EPI runs were then co-registered to the mean EPI image.

For the general linear model (GLM), regressors were defined for each sequence separately for both preparation and production. Preparation- and production-related BOLD responses were independently modelled from ‘No-Go’ and ‘Go’ trials, respectively, to tease out activity from these brief trial phases despite given the haemodynamic response lag ^64^. The preparation regressor consisted of boxcar function starting at the moment of the Sequence cue in ‘No-Go’ trials and lasting for the duration of the maximum possible preparation phase (2500 ms). The production regressor consisted of a boxcar function starting at the onset of the first press with a fixed duration of 0 (constant-duration impulse), to capture activity related to sequence initiation and extract sequence production related activity from the first finger press that was matched across sequences within each participant. Additionally, we included regressors or no interest: 1) Error trials (incorrect or premature presses during ‘Go’ trials and presses during No-Go trials) which were modelled from sequence cue onset to the end of the ITI, 2) the preparation period in ‘Go’ trials (1000-2500 ms from Sequence cue) and 3) the temporal derivate of each regressor. The boxcar model was then convolved with the standard hemodynamic response function. To remove the influence of movement-related artifacts, we used a weighted least-squares approach ^65^. The GLM regressors were optimised based on independent pilot data (N=10).

### surface reconstruction

Cortical surface reconstruction was conducted on each participant’s T1 anatomical image using Freesurfer’s recon-all function ^66^. Surface structures were then co-registered to the symmetrical Freesurfer average atlas ^67^ using surface Caret ^68^. Searchlights for multivariate Pattern Analysis (MVPA) were then defined on each individual surface using the node maps provided by the surface reconstruction and displayed in atlas space.

### Cross-sectional and region of interest analysis

Two cross-sections were defined upon the cortical surface: 1) anterior to posterior, running from PMd to OPJ and 2) ventral to dorsal, running from PMv to SMA. These cross-sections were taken from a previous study ^12^. Data points along these axes were extracted to provide a continuous measure along the cortical surface, which was then subjected to a non-parametric permutation analysis to identify clusters which were significantly above baseline^29^. This was conducted as a one-tailed test, with 10,000 permutations.

Region of interest analysis was conducted using the Caret toolbox ^68^ on RoIs which were defined based on several previous studies ^12,20,21^, consisting of PMv, PMd, M1, S1, SMA/pre-SMA, SPCa, and SPCp. Z-values for each classifier were averaged within regions to give an overall value for each decoder, these values were calculated from unsmoothed individual data. One-sample t-tests against chance level (zero) then identified significantly above-chance decoding values on these cross-sections.

### Multi-variate pattern analysis of fMRI

Multivariate pattern analysis (MVPA) was conducted using a custom-written MATLAB code to detect sequence-specific representations ^12,14^. We used a searchlight of 160 voxels and a maximum searchlight radius of six millimetres. Each searchlight was run on each individual’s cortical surface-reconstructed anatomy, projected onto the Freesurfer average atlas ^67^. The classification accuracy for each searchlight (cf. classification procedures below) was assigned to the centre of each searchlight. A classification accuracy map was generated by moving the searchlight across the cortical surface ^30^. Mean patterns and common voxel-by-voxel co-variance matrices were extracted for each class from training data set (five of the six imaging runs), and then a gaussian linear discriminant classifier was used to distinguish between the same classes in a test data set (the remaining imaging run). For overall sequence classification (Supplementary Figure S1b), the classifier was trained to distinguish between sequences on five runs and then tested on the remaining run, performed on betas estimated from the sequence preparation and production periods independently. Discrimination was then cross-validated across runs (six cross-validation folds).

The factorised classification of finger order, timing, and integrated order and timing followed the previous approach ^12^. For the decoding of sequence timing, the classifier was trained to distinguish between two sequences with differing timing but matching order across five runs and was then tested on two sequences with the same two timings paired with a different order in the remaining run. This classification was then cross-validated across runs and across training/test sequences, for a total of 12 cross-validation folds. For the decoding of sequence order, the classifier was trained to distinguish between two sequences of differing orders paired with the same timing and tested on two sequences with the same two orders when paired with a different timing and underwent the same cross-validation procedure. The integrated classifier was analogous to overall sequence decoding, but here the mean activity for each timing (collapsed across two orders) and finger order (collapsed across two timings) condition within each run was subtracted from the overall activity for each run, separately. This allowed for the measurement of residual activity patterns that were not explained by a linear combination of timing and order. For better comparability across classifiers and for the group analysis, the classification accuracies were transformed to z-scores, assuming a binomial distribution of the number of correct guesses. We then tested these z-scores against zero (chance level) on cortical cross-sections of interest and in pre-defined ROIs across participants for statistical analysis.

## Supporting information

Supplementary Materials Yewbrey_etal_bioRxiv_2022

## Acknowledgements

The authors wish to thank David McKiernan for the construction and technical support of the force keyboard device, and Prof Paul Downing, Prof Paul Mullins, and Dr Ken Valyear for their useful comments on the study design and MRI acquisition. This work was supported by the Academy of Medical Sciences Springboard Award (SBF006\1052, K.K.).

