## Supplementary Materials Yewbrey_etal_bioRxiv_2022 for "Cortical patterns shift from sequence feature separation during planning to integration during motor execution"

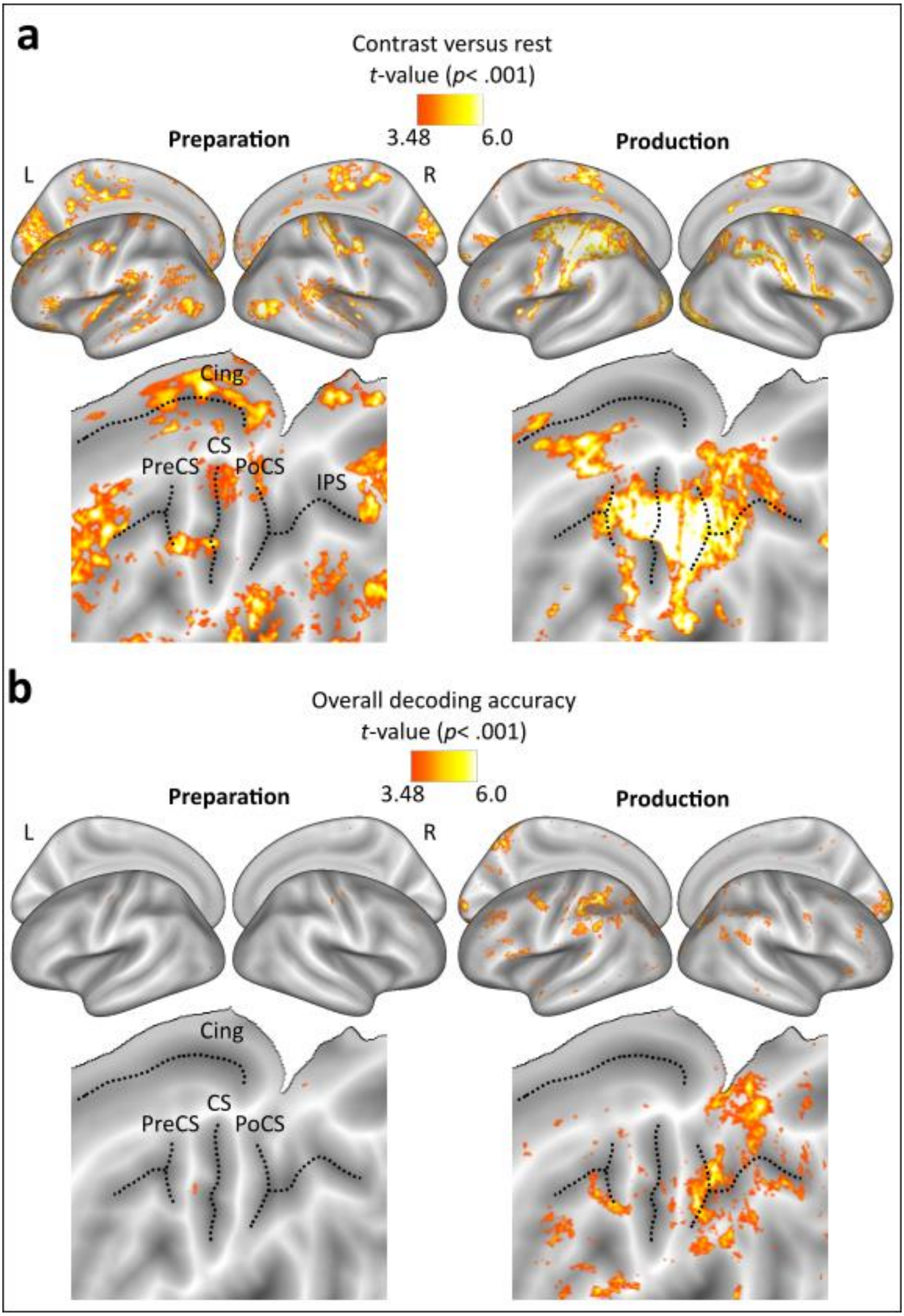

**Figure S1** T-value maps of contrasts versus rest and overall sequence decoding accuracy versus chance. (a) Inflated surface maps are shown in upper panels and flat maps in lower panels, displaying contrasts of preparation versus rest on the left and production versus rest on the right panels, respectively. (b) As above but displaying overall sequence decoding using the MVPA to decode between all four target sequences.

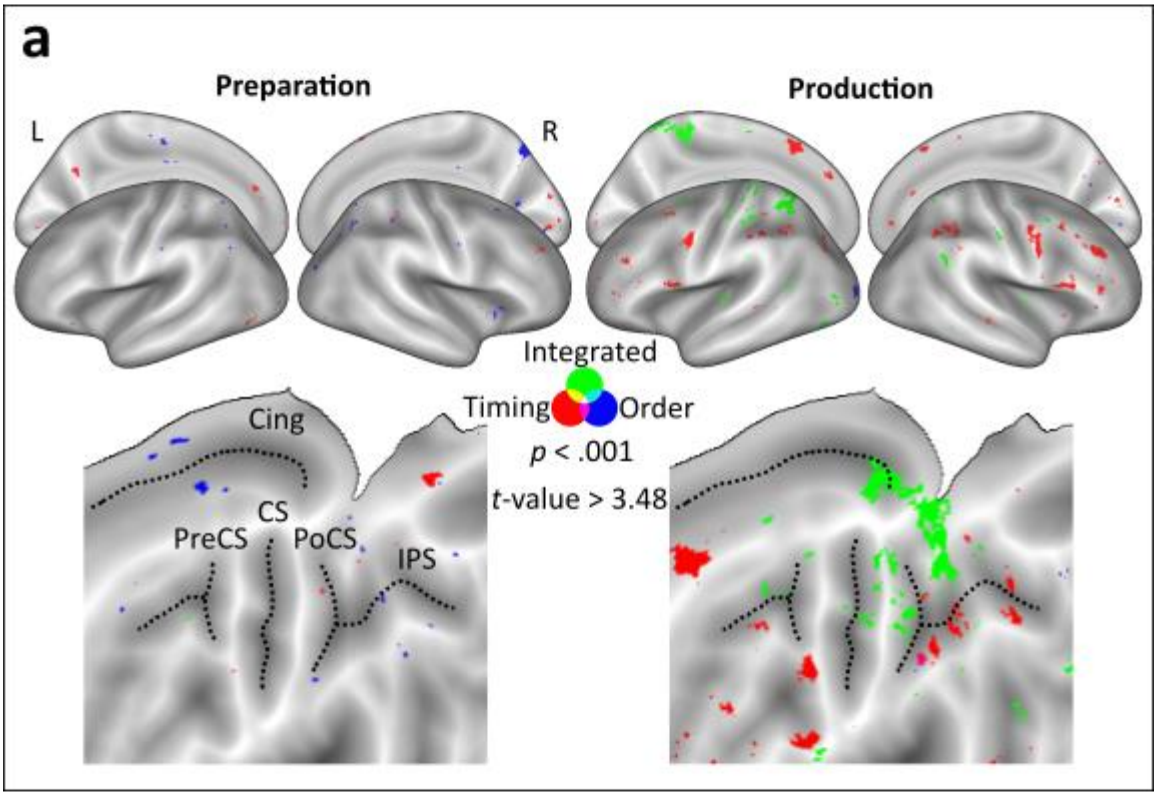

**Figure S2.** T-value maps of whole-brain sequence feature decoding. (a) Inflated surface maps are shown in upper panels and flat maps in lower panels, displaying order, timing, and integration decoding during preparation on the left and production on the right panels, respectively.

**Table S1**

*Distribution of trial types across experimental phases. The first half of training 2 occurred on Day 1, the second half on Day 2.*

| | **Day 1** | | | **Day 2** | | | **Day 3** | |
|---|---|---|---|---|---|---|---|---|
| | Example | Pre-Training Probe | Training 1 | Training 2 | Training 3 | Post-Training Probe | Refresher | Test (fMRI) |
| **Instructed Trials** | 4 (33%) | 32 (100%) | 16 (80%) | 8 (40%) | 0 | 32 (100%) | 8 (40%) | 0 |
| **Memory Trials** | 4 (33%) | 0 | 0 | 8 (40%) | 16 (80%) | 0 | 8 (40%) | 24 (50%) |
| **No-go Trials** | 4 (33%) | 0 | 4 (20%) | 4 (20%) | 4 (20%) | 0 | 4 (20%) | 24 (50%) |
| **Total Trials per block** | 12 | 32 | 20 | 20 | 20 | 32 | 20 | 48 |
| **Number of blocks** | 1 | 4 | 12 | 12 | 12 | 4 | 2 | 6 |
| **Repetitions per condition per block** | 2 | 8 | 4 | 4 | 4 | 8 | 4 | 6 |

**Table S2**

*Significant whole-brain clusters for % signal change contrasts against rest and MVPA classification accuracy against baseline. Results of surface-based random effects analysis (N = 24) with an uncorrected threshold of t(23) > 3.48, p<0.001. p (cluster) is the cluster-wise p-value for the cluster of that size. The p-value is corrected over the cortical surface using the area of the cluster* (*Worsley et al., 1996*). *The cluster coordinates reflect the location of the cluster peak in MNI space.*

| | | | | | MNI | | |
|---|---|---|---|---|---|---|---|
| Contrasts | Area (Brodmann Area) | Extent | *p* (cluster) | Peak *t* | X | Y | Z |
| Preparation vs. Rest | **Contralateral** | | | | | | |
| | Extrastriate Vis Cortex (BA18) | 4886.16 | <.001 | 7.21 | -12 | -60 | -5 |
| | Pre-SMA (BA32) | 2249.6 | <.001 | 6.99 | -11 | 14 | 50 |
| | Primary Auditory Cortex (BA41) | 940.58 | <.001 | 7.84 | -44 | -29 | 11 |
| | Posterior Cingulate (BA23) | 733.45 | <.001 | 6.86 | -11 | -38 | 31 |
| | Anterior Insula (BA48) | 706.11 | <.001 | 6.55 | -37 | -12 | 3 |
| | Occipitotemporal Area (BA37) | 698.86 | <.001 | 6.39 | -38 | -62 | 2 |
| | Anterior Insula (BA48) | 523.72 | <.001 | 6.67 | -46 | 18 | 11 |
| | M1 (BA4) | 483.3 | <.001 | 6.86 | -46 | -10 | 44 |
| | Inferior Parietal (BA39) | 480.37 | <.001 | 5.48 | -50 | -59 | 26 |
| | Extrastriate Vis Cortex (BA18) | 477.2 | <.001 | 6.19 | -20 | -90 | 6 |
| | S1 (BA2) | 429.79 | <.001 | 5.06 | -18 | -39 | 59 |
| | Orbitofrontal (BA47) | 373.43 | <.001 | 6.82 | -23 | 29 | 2 |
| | Superior Parietal (BA7) | 293.49 | <.001 | 5.52 | -21 | -49 | 51 |
| | Posterior Cingulate (BA23) | 274.22 | <.001 | 6.71 | -13 | -55 | 26 |
| | Pre-SMA (BA32) | 231.75 | <.001 | 5.46 | -9 | 46 | 6 |
| | Middle Temporal (BA21) | 179.26 | <.001 | 6.06 | -46 | -37 | -3 |
| | Middle Temporal (BA21) | 125.41 | <.001 | 6.18 | -54 | -26 | -7 |
| | Inferior Parietal (BA39) | 122.42 | <.001 | 5.42 | -50 | -71 | 16 |
| | Anterior Insula (BA48) | 115.35 | <.001 | 4.46 | -28 | 31 | 29 |
| | Extrastriate Vis Cortex (BA19) | 98.81 | <.001 | 5.38 | -19 | -53 | 3 |
| | Wernicke's Area (BA22) | 92.06 | <.001 | 4.82 | -53 | -19 | 3 |
| | Occipitotemporal Area (BA37) | 86.6 | <.001 | 4.75 | -43 | -41 | -19 |
| | Anterior Prefrontal (BA10) | 54.95 | 0.007 | 4.49 | -6 | 56 | 13 |
| | Wernicke's Area (BA22) | 53.73 | 0.008 | 4.5 | -57 | -37 | 3 |
| | Anterior Insula (BA48) | 53.07 | 0.009 | 4.35 | -54 | -1 | 11 |
| | Posterior Cingulate (BA23) | 51.92 | 0.01 | 6.21 | -4 | -26 | 38 |
| | Anterior Insula (BA48) | 51.89 | 0.01 | 5.99 | -55 | 3 | 14 |
| | Primary Auditory Cortex (BA42) | 48.98 | 0.014 | 4.71 | -57 | -37 | 16 |
| | Superior Parietal (BA5) | 39.88 | 0.043 | 5 | -12 | -44 | 45 |
| | **Ipsilateral** | | | | | | |
| | Extrastriate Vis Cortex (BA18) | 2605.25 | <.001 | 7.71 | -14 | -73 | -8 |
| | Extrastriate Vis Cortex (BA19) | 1523.99 | <.001 | 7.2 | -21 | -83 | 16 |

| | | | | | | | |
|---|---|---|---|---|---|---|---|
| | S1 (BA3) | 1498.75 | <.001 | 7.89 | -37 | -26 | 37 |
| | Superior Parietal (BA5) | 1142.07 | <.001 | 7.88 | -17 | -53 | 47 |
| | Occipitotemporal Area (BA37) | 883.28 | <.001 | 7.65 | -46 | -63 | -1 |
| | Ventral Temporal (BA20) | 671.28 | <.001 | 6.27 | -49 | -37 | 14 |
| | Superior Parietal (BA40) | 336.82 | <.001 | 5.56 | -34 | -52 | 49 |
| | Anterior Cingulate (BA24) | 282.58 | <.001 | 5.86 | -6 | 31 | 12 |
| | Extrastriate Vis Cortex (BA18) | 274.65 | <.001 | 4.87 | -23 | -87 | 3 |
| | Anterior Insula (BA48) | 268.99 | <.001 | 6.19 | -41 | -24 | 20 |
| | Superior Parietal (BA40) | 245.49 | <.001 | 5.49 | -31 | 36 | 35 |
| | Anterior Insula (BA48) | 159.52 | <.001 | 6.36 | -55 | -8 | 13 |
| | Pre-SMA (BA32) | 145.62 | <.001 | 5.05 | -10 | 54 | 22 |
| | Middle Temporal (BA21) | 122.81 | <.001 | 4.83 | -53 | -50 | 3 |
| | Pre-SMA (BA32) | 106.49 | <.001 | 5.86 | -5 | 44 | 7 |
| | Anterior Cingulate (BA24) | 93.31 | <.001 | 4.94 | -9 | 11 | 30 |
| | Subgenual Area (BA25) | 56.97 | 0.003 | 5.21 | -4 | 22 | 6 |
| | Posterior Cingulate (BA23) | 56.86 | 0.003 | 5.13 | -4 | -12 | 34 |
| | S1 (BA3) | 48.96 | 0.008 | 5.22 | -54 | -17 | 37 |
| | Middle Temporal (BA21) | 48.93 | 0.008 | 4.64 | -49 | -34 | -4 |
| | Posterior Cingulate (BA23) | 44.43 | 0.015 | 5.35 | -4 | -30 | 32 |
| | Pre-SMA (BA32) | 43.95 | 0.016 | 4.43 | -6 | 48 | 27 |
| | Dorsolateral Prefrontal (BA9) | 40.09 | 0.026 | 4.57 | -12 | 34 | 46 |
| | Ectosplenial Area (BA26) | 36.16 | 0.045 | 4.43 | -6 | -45 | 25 |
| Production vs. Rest | **Contralateral** | | | | | | |
| | Superior Parietal (BA40) | 9012.24 | <.001 | 13.05 | -34 | -34 | 38 |
| | Extrastriate Vis Cortex (BA18) | 1166.68 | <.001 | 7.05 | -30 | -86 | -12 |
| | SMA (BA6) | 741.62 | <.001 | 9.5 | -3 | -16 | 59 |
| | M1 (BA4) | 717.53 | <.001 | 8.91 | -53 | -1 | 30 |
| | Extrastriate Vis Cortex (BA18) | 701.31 | <.001 | 6.37 | -18 | -71 | -2 |
| | Extrastriate Vis Cortex (BA18) | 561.59 | <.001 | 7.03 | -20 | -86 | -21 |
| | Anterior Insula (BA48) | 400.9 | <.001 | 6.84 | -33 | 2 | 7 |
| | Extrastriate Vis Cortex (BA18) | 174.37 | <.001 | 5.88 | -15 | -92 | -12 |
| | Posterior Cingulate (BA23) | 103.37 | <.001 | 5.43 | -5 | -5 | 38 |
| | Anterior Insula (BA48) | 86.36 | <.001 | 7.69 | -26 | 13 | 16 |
| | Primary Auditory Cortex (BA41) | 79.07 | <.001 | 5.6 | -48 | -44 | 25 |
| | Broca's Area (BA45) | 77.51 | <.001 | 5.8 | -45 | 30 | 32 |
| | Extrastriate Vis Cortex (BA18) | 45.13 | 0.001 | 4.98 | -37 | -69 | -16 |
| | **Ipsilateral** | | | | | | |
| | Anterior Insula (BA48) | 2828.07 | <.001 | 9.24 | -51 | -42 | 34 |
| | Extrastriate Vis Cortex (BA19) | 2365.33 | <.001 | 7.46 | -46 | -73 | -21 |
| | Extrastriate Vis Cortex (BA18) | 1295.15 | <.001 | 6.91 | -9 | -81 | 32 |
| | M1 (BA4) | 974.65 | <.001 | 7.87 | -55 | -4 | 35 |
| | V1 (BA17) | 542.45 | <.001 | 5.52 | -9 | -71 | 5 |

| | | | | | | | |
|---|---|---|---|---|---|---|---|
| | SMA (Medial BA6) | 513.52 | <.001 | 8.01 | -4 | -14 | 61 |
| | PMd (Dorsal BA6) | 285.07 | <.001 | 7.48 | -29 | -7 | 48 |
| | Extrastriate Vis Cortex (BA18) | 266.94 | <.001 | 6.91 | -21 | -89 | -13 |
| | M1 (BA4) | 239.01 | <.001 | 8.4 | -33 | -21 | 56 |
| | Posterior Cingulate (BA23) | 159.35 | <.001 | 6.4 | -8 | -24 | 24 |
| | Anterior Cingulate (BA24) | 70.39 | <.001 | 5.71 | -9 | 11 | 31 |
| | Pars Opercularis (BA44) | 62.97 | <.001 | 5.49 | -48 | 19 | 33 |
| | Pars Triangularis (BA45) | 42.9 | 0.008 | 5.16 | -43 | 34 | 32 |
| | Pars Triangularis (BA45) | 39.1 | 0.014 | 4.87 | -38 | 40 | 18 |

23

| | | | | | MNI | | |
|---|---|---|---|---|---|---|---|
| Above-chance classification accuracy | Area (Brodmann Atlas) | Extent | *p* (cluster) | Peak *t* | X | Y | Z |
| Overall Preparation | | | | | | | |
| Overall Production | **Contralateral** | | | | | | |
| | Superior Parietal (BA40) | 1168.72 | <.001 | 6.69 | -32 | -50 | 37 |
| | Extrastriate Vis Cortex (BA18) | 1121.03 | <.001 | 6.41 | -8 | -79 | 34 |
| | SMA (BA6) | 354.11 | <.001 | 5.5 | -41 | -9 | 44 |
| | Broca's Area (BA44) | 344.4 | <.001 | 6.02 | -54 | 13 | 26 |
| | Extrastriate Vis Cortex (BA19) | 339.11 | <.001 | 5.66 | -29 | -80 | 21 |
| | Anterior Insula (BA48) | 253.22 | <.001 | 5.03 | -56 | -34 | 28 |
| | Broca's Area (BA44) | 245.75 | <.001 | 5.17 | -36 | 21 | 33 |
| | Inferior Parietal (BA39) | 244.39 | <.001 | 5.16 | -41 | -65 | 33 |
| | Extrastriate Vis Cortex (BA18) | 232.48 | <.001 | 5.62 | -16 | -87 | -7 |
| | Anterior Insula (BA48) | 180.79 | <.001 | 5.66 | -55 | -45 | 29 |
| | Extrastriate Vis Cortex (BA18) | 165.59 | <.001 | 4.81 | -13 | -88 | -20 |
| | Anterior Insula (BA48) | 72.9 | 0.002 | 4.91 | -36 | 12 | 27 |
| | Anterior Insula (BA48) | 65.59 | 0.006 | 4.48 | -43 | 28 | 17 |
| | Superior Parietal (BA7) | 62.01 | 0.008 | 4.41 | -23 | -64 | 37 |
| | Extrastriate Vis Cortex (BA18) | 61.88 | 0.008 | 4.24 | -27 | -89 | 7 |
| | Temporal Pole (BA38) | 48.8 | 0.032 | 5.63 | -55 | 10 | -4 |
| | S1 (BA2) | 46.31 | 0.044 | 4.22 | -49 | -27 | 40 |
| | Frontal Eye Fields (BA8) | 44.85 | 0.051 | 4.4 | -22 | 19 | 42 |
| | **Ipsilateral** | | | | | | |
| | V1 (BA17) | 1072.6 | <.001 | 8.19 | -17 | -99 | -5 |
| | Extrastriate Vis Cortex (BA19) | 214.86 | <.001 | 8.05 | -28 | -75 | 6 |
| | Middle Temporal (BA21) | 187.47 | <.001 | 5.28 | -57 | -54 | 17 |
| | Extrastriate Vis Cortex (BA19) | 175.75 | <.001 | 4.84 | -31 | -79 | 29 |
| | Occipitotemporal Area (BA37) | 135.18 | <.001 | 5.06 | -39 | -62 | 15 |
| | Extrastriate Vis Cortex (BA18) | 125.11 | <.001 | 4.61 | -13 | -77 | 32 |
| | Superior Parietal (BA40) | 104.19 | <.001 | 4.53 | -44 | -42 | 37 |
| | Pars Opercularis (BA44) | 86.9 | <.001 | 4.98 | -48 | 24 | 29 |

| | | | | | | | |
|---|---|---|---|---|---|---|---|
| | Superior Parietal (BA7) | 80.06 | <.001 | 4.97 | -20 | -70 | 45 |
| | Superior Parietal (BA40) | 72.6 | 0.002 | 4.81 | -45 | -47 | 32 |
| | S1 (BA3) | 71.55 | 0.002 | 4.63 | -52 | -20 | 38 |
| | Pars Triangularis (BA45) | 57.6 | 0.008 | 4.8 | -41 | 31 | 8 |
| | S1 (BA2) | 46.3 | 0.032 | 4.74 | -58 | -31 | 31 |

24

| | | | | | MNI | | |
|---|---|---|---|---|---|---|---|
| Above-chance classification accuracy | Area (Brodmann Atlas) | Extent | *p* (cluster) | Peak *t* | X | Y | Z |
| Integrated Preparation | | | | | | | |
| Integrated Production | **Contralateral** | | | | | | |
| | Superior Parietal (BA7) | 606.08 | <.001 | 7.27 | -17 | -63 | 56 |
| | Superior Parietal (BA5) | 183.08 | <.001 | 6.6 | -14 | -47 | 46 |
| | Extrastriate Vis Cortex (BA19) | 62.85 | 0.008 | 4.67 | -47 | -66 | -19 |
| | S1 (BA2) | 46.26 | 0.05 | 4.34 | -28 | -44 | 58 |
| | **Ipsilateral** | | | | | | |
| | Inferior Parietal (BA39) | 69.53 | 0.004 | 5.81 | -51 | -54 | 29 |
| | Superior Parietal (BA7) | 63.4 | 0.008 | 5.01 | -16 | -65 | 49 |
| Order Preparation | **Contralateral** | | | | | | |
| | Extrastriate Vis Cortex (BA18) | 61.68 | 0.012 | 4.33 | -23 | -73 | -18 |
| | Extrastriate Vis Cortex (BA18) | 56.03 | 0.022 | 5.04 | -23 | -82 | -13 |
| | **Ipsilateral** | | | | | | |
| | Extrastriate Vis Cortex (BA18) | 199.36 | <.001 | 5.56 | -9 | -80 | 30 |
| | Extrastriate Vis Cortex (BA19) | 80.09 | 0.004 | 4.9 | -21 | -62 | -5 |
| Order Production | **Contralateral** | | | | | | |
| | Extrastriate Vis Cortex (BA18) | 118.04 | <.001 | 6.51 | -24 | -94 | -10 |
| Timing Preparation | **Ipsilateral** | | | | | | |
| | Extrastriate Vis Cortex (BA18) | 58.41 | 0.026 | 4.69 | -19 | -81 | -22 |
| Timing Production | **Contralateral** | | | | | | |
| | SMA (BA6) | 130.73 | <.001 | 5.13 | -3 | 3 | 53 |
| | Broca's Area (BA44) | 128.93 | <.001 | 4.94 | -49 | 9 | 13 |
| | S1 (BA3) | 114.82 | <.001 | 5.78 | -50 | -14 | 39 |
| | Superior Parietal (BA40) | 76.95 | 0.01 | 5.57 | -40 | -45 | 36 |
| | Extrastriate Vis Cortex (BA19) | 59.21 | 0.044 | 4.36 | -35 | -81 | 16 |
| | Inferior Parietal (BA39) | 57.46 | 0.05 | 4.25 | -35 | -57 | 28 |

| | | | | | | | |
|---|---|---|---|---|---|---|---|
| | **Ipsilateral** | | | | | | |
| | M1 (BA4) | 199.04 | <.001 | 6.08 | -52 | -11 | 40 |
| | PMv (Ventral BA6) | 164.2 | <.001 | 6.26 | -55 | 7 | 23 |
| | Inferior Parietal (BA39) | 155.11 | <.001 | 5.76 | -39 | -48 | 27 |
| | Pars Opercularis (BA44) | 151.26 | <.001 | 5.42 | -41 | 22 | 35 |
| | Inferior Parietal (BA39) | 93.54 | 0.002 | 5.14 | -35 | -60 | 28 |
| | Pars Triangularis (BA45) | 59.82 | 0.034 | 4.21 | -43 | 27 | 17 |
